# Identifying HRV and EEG correlates of well-being using ultra-short, portable, and low-cost measurements

**DOI:** 10.1101/2024.02.23.581823

**Authors:** Cédric Cannard, Arnaud Delorme, Helané Wahbeh

## Abstract

Wearable electroencephalography (EEG) and electrocardiography (ECG) devices may offer a non-invasive, user-friendly, and cost-effective approach for assessing well-being (WB) in real-world settings. However, challenges remain in dealing with signal artifacts (such as environmental noise and movements) and identifying robust biomarkers. We evaluated the feasibility of using portable hardware to identify potential EEG and heart-rate variability (HRV) correlates of WB. We collected simultaneous ultrashort (2-minute) EEG and ECG data from 60 individuals in real-world settings using a wrist ECG electrode connected to a 4-channel wearable EEG headset. These data were processed, assessed for signal quality, and analyzed using the open-source EEGLAB BrainBeats plugin to extract several theory-driven metrics as potential correlates of WB. Namely, the individual alpha frequency (IAF), frontal and posterior alpha asymmetry, and signal entropy for EEG. SDNN, the low/high frequency (LF/HF) ratio, the Poincaré SD1/SD2 ratio, and signal entropy for HRV. We assessed potential associations between these features and the main WB dimensions (hedonic, eudaimonic, global, physical, and social) implementing a pairwise correlation approach, robust Spearman’s correlations, and corrections for multiple comparisons. Only 8 files showed poor signal quality and were excluded from the analysis. Eudaimonic (psychological) WB was positively correlated with SDNN and the LF/HF ratio. EEG posterior alpha asymmetry was positively correlated with Physical WB (i.e., sleep and pain levels). No relationships were found with the other metrics, or between EEG and HRV metrics. These physiological metrics enable a quick, objective assessment of well-being in real-world settings using scalable, user-friendly tools.

## Introduction

Well-being (WB), a concept rooted in ancient Greek philosophy (Aristotle, 2000), is generally understood through two primary dimensions: the hedonic, which concentrates on the balance of positive and negative affect (i.e., affective WB) Kahneman et al., 1999; Ryff et al., 2008), and the eudaimonic (i.e., psychological WB), focusing on self-realization, life purpose, life satisfaction, and the actualization of one’s potential (Deci and Ryan, 2008; Kahneman et al., 1999; Ryff, 2014). However, it is a multifaceted concept influenced by various factors (e.g., physical condition, social relationships, spiritual beliefs, culture, socioeconomic status).

Measuring WB plays a crucial role not only in treating patients but also in preventing mental illness in healthy individuals and improving their overall happiness and life function (Keyes et al., 2010; Ruini and Fava, 2009). Studies indicate that lower levels of WB are linked with an increased risk for cardiovascular disease and mortality (Chalmers et al., 2014), while higher levels are associated with resilience and the ability to recover from stressors and negative experiences (Fredrickson, 2004). Indeed, stress is a key disruptor of WB, affecting the balance between the sympathetic (SNS) and parasympathetic (PNS) branches of the autonomic nervous system (ANS) and activating the hypothalamic-pituitary-adrenal axis (Tsigos and Chrousos, 2002). While acute stress can be beneficial by increasing lucidity, performance, and chances of survival, chronic stress results in cognitive and emotional disruptions, including reduced cognitive flexibility and increased negative bias and mental disorders (e.g., depression and anxiety). These changes are largely influenced by individual perceptions of stress, stress responses, and emotion regulation strategies (Aldao et al., 2010; Edwards and Pinna, 2020).

WB is often assessed through subjective questionnaires (Ryff and Singer, 1996). However, the reliability of these subjective measures has been questioned, leading to suggestions that physiological measures or biomarkers might offer a more objective and reliable assessment of WB levels. These biomarkers could help us understand the underlying mechanisms of WB and modulate WB-related physiological processes, potentially through biofeedback interventions (Millear et al., 2008; Ruini et al., 2009). The utilization of physiological tools for early detection of cognitive and emotional activity disruptions is a significant development in their prevention, reducing costs for individuals and society. These tools are vital for measuring psychological and physiological changes, tracking the effectiveness of interventions and treatments, and detecting relapses. Identifying multimodal markers is crucial for understanding neurophysiological system interactions and early detection of cardiovascular disease or mental disorder signs before symptoms become severe. Integrating objective physiological measures with subjective WB reports could provide a more comprehensive understanding of WB. In some cases, these physiological markers could serve as standalone indicators, especially where subjective reports are unavailable or unreliable.

Electrocardiography (ECG) and electroencephalography (EEG) are promising tools for examining the complex bidirectional dynamics between the autonomous nervous system (ANS) and the central nervous system (CNS). These tools are non-invasive, mobile, and affordable, and allow easy data collection in real-world settings with recent advancements in wearable technologies. The Muse headset (InteraXon Inc.) has been validated for collecting EEG signals comparable to state-of-the-art systems (Cannard et al., 2021a; Krigolson et al., 2017), including relevant EEG metrics such as the alpha asymmetry and the individual alpha frequency (IAF), described in more detail below. The Muse has been used in many recent EEG studies (e.g., Amores et al., 2018; Arsalan et al., 2019; Asif et al., 2019; Hashemi et al., 2016). Similarly, ECG recordings from the wrists can capture the QRS complex reliably with proper signal-denoising tools (Bent et al., 2020; Lynn et al., 2013). Despite the inherent ‘noisiness’ in data from such low-cost devices, these signals can offer valuable insights into the physiological correlates of WB (Cannard et al., 2021b). Hence, for this study, we built an ECG electrode that connects to the Muse auxiliary port, allowing us to collect simultaneous EEG-ECG signals easily and affordably in real-world settings. This study explores integrating wearable technology to objectively measure WB, specifically a Muse headband with four EEG channels and a custom-built ECG electrode connected to the participants’ wrists.

Heart-rate variability (HRV; the time variation between heartbeats), derived from ECG signal, is a simple and recognized measure of cardiovascular regulation and physiological adaptation mediated by the ANS and shows great promise as a correlate of WB. While 24-hour ECG recordings are the gold standard for capturing slow regulatory mechanisms and allow medical prognostic, short (15 min) and ultrashort (<5 min) measurements also provide valuable insights into the PNS activity and can be easily collected with wearable technologies (Shaffer et al., 2020; Shaffer and Ginsberg, 2017). Short-HRV is positively correlated with positive dimensions of WB, such as emotion regulation, adaptability, resilience, and executive functioning (Appelhans and Luecken, 2006; Shaffer et al., 2014) and negatively correlated with negative dimensions of WB, such as worry, mental rumination, negative affect, and mental disorders (Beauchaine and Thayer, 2015; Chalmers et al., 2014).

A popular time-domain HRV measure to capture the balance between the SNS and PNS branches of the ANS is the standard deviation of the normal-to-normal (NN) intervals (SDNN) and is therefore included in this study as a potential correlate of WB. A higher SDNN score is often interpreted as a sign of a healthy, adaptable ANS, which allows the body to efficiently manage stress, recover from stressors, and maintain homeostasis, contributing to improved physical health, reduced stress levels, and enhanced emotion al regulation (Kemp and Quintana, 2013; Thayer et al., 2012). In the frequency domain, low-(LF; 0.04–0.15 Hz) and high-frequency (HF; 0.15 - 0.40 Hz) power and the LF/HF ratio are promising correlates of WB, with several direct associations reported in the literature (Boman, 2018; Sloan et al., 2017; Trimmel, 2015). In addition, nonlinear HRV measures (e.g., signal entropy) show promise at providing insights into complex, bidirectional interactions between various physiological systems that may not be picked up by frequency-domain measures (Shaffer and Ginsberg, 2017; Stein and Reddy, 2005). A popular nonlinear measure of HRV is the Poincaré plot, a geometric representation of the correlation between consecutive heartbeats, thought to indicate the balance between SNS and PNS activity. Alterations in Poincaré plot metrics have been linked to stress-related autonomic dysregulation (Shaffer and Ginsberg, 2017). Similarly, low entropy values (another popular nonlinear RHV metric) are associated with stress, anxiety, and depression, and are suspected to reflect dysregulations in the ANS (Chalmers et al., 2014; Shaffer and Ginsberg, 2017).

A promising EEG correlate of WB that can be collected with these devices is EEG alpha asymmetry. EEG alpha asymmetry is the difference in alpha power (8-13 Hz) between the brain’s right and left hemispheres and is a promising measure of the brain’s motivational and emotional systems (Allen and Reznik, 2015; Coan and Allen, 2004; Harmon-Jones et al., 2010). Greater left than right frontal alpha power is typically associated with difficulties in disengaging from negative or avoidance-motivated information (leading to depression and anxiety). In contrast, greater right than left frontal alpha power is linked to challenges in inhibiting positive or approach-motivated distractions (leading to addiction and risk-taking behaviors). Urry and colleagues found that higher levels of eudaimonic and hedonic WB corresponded with greater left than right frontal alpha power in response to emotional stimuli, underscoring its potential as a direct indicator of WB (Urry et al., 2004). More recently, we found that greater left-than-right alpha power in the posterior regions was positively associated with multidimensional WB in 230 participants using a wearable headset (Cannard et al., 2021b). Another potentially promising EEG metric is the individual alpha frequency (IAF; referring to the peak frequency within the alpha band). Higher IAF is generally associated with better cognitive abilities, such as memory and problem-solving skills, which may contribute to effective stress management and emotional regulation, thus potentially enhancing well-being (Klimesch, 1999). In addition to alpha asymmetry and IAF, entropy measures from EEG signals (e.g., entropy) provide insights into the complexity and irregularity of brain activity, offering a different perspective on neural dynamics relative to traditional frequency-domain information. Variations in entropy could reflect differences in psychological or emotional brain states (Hosseini and Naghibi-Sistani, 2011; Thul et al., 2016). This approach highlights the potential of using entropy as a biomarker for mental health and WB, offering a quantitative measure that could complement traditional psychological assessments.

In summary, for their reliability in ultrashort recordings and their potential to capture physiological activity related to factors contributing to WB, our study focuses on the following EEG and HRV metrics:

- EEG: individual alpha frequency (IAF), alpha asymmetry, and signal entropy.
- HRV: SDNN, low/high-frequency ratio (LF/HF ratio), the Poincare SD1/SD2 ratio, and signal entropy

In addition to examining ECG and EEG metrics independently, some studies suggest that EEG and HRV measures may present some relationships, which could inform on potential dynamics between the neural and cardiovascular systems. For instance, EEG delta power was negatively correlated with LF-HRV power and the LF/HF ratio during sleep (Abdullah et al., 2010; Ako et al., 2003). Hence, we conduct an additional exploratory analysis to evaluate whether these EEG and HRV measures may be correlated in our sample at rest.

Thus, with the overall goal of evaluating the relevant physiological correlates of WB described above with a low-cost portable mobile device, the objectives of this study were to:

1. Evaluate the feasibility of collecting ECG and EEG signals with a low-cost portable system with satisfying quality to extract relevant information.
2. Assess whether these EEG metrics are associated with various dimensions of WB, and if so, which ones.
3. Assess whether these HRV metrics are associated with various dimensions of WB, and if so, which ones.
4. Explore whether EEG and HRV metrics show relationships, potentially informing on brain-heart interplay.

## Methods

### Participants

Sixty participants (73.2% women; age range: 31-73 years old; mean age: 54; SD age = 12) were recruited through the Institute of Noetic Sciences (IONS)’ Discovery Laboratory research program (Cannard et al., 2021b; Wahbeh et al., 2022b, 2022a). Excluded from the study were people younger than 18 years of age, unable to read or understand the consent form, and acute or chronic illness precluding completion of measurements. Upon arrival at the research laboratory, participants were briefly interv iewed by the research assistants to ensure they met the inclusion/exclusion criteria and were then directed to a desk where the equipment was available. Participants volunteered and were not compensated for their participation. All questionnaires were optional and anonymous, and all study activities were approved by the IONS institutional review board (IORG#0003743).

### Well-being data collection

Participants completed a battery of validated questionnaires in SurveyMonkey to capture the main dimensions of WB:

1) **Global WB** using the Arizona integrative outcome scale (AIOS) assessing the global sense of physical, social, psychological, affective, and spiritual WB over the past 24 hours (Bell et al., 2004; Otto et al., 2010; Tuason et al., 2021).
2) **Eudaimonic WB** using the average score from items of the Cloninger Scale (Cloninger et al., 1997), capturing the commitment to making the world a better place (life purpose) and the sense of alignment between natural responses and long-term goals and principles.
3) **Hedonic WB** using the positive and negative affect schedule short form (I-PANAS-SF (Thompson, 2007) indicating to what extent participants have felt 10 predefined emotions during the past few days (5 positive items being alert, inspired, determined, attentive, active and 5 negative items being upset, hostile, ashamed, nervous, afraid). The positive-to-negative-affect ratio was calculated by dividing the summed positive scores by the summed negative scores. A positive ratio indicates more positive than negative affect and vice versa.
4) **Physical WB** using the summed sleep disturbance (Sleep Quality Scale; Cappelleri et al., 2009) and pain intensity (Bijur et al., 2001; Boonstra et al., 2008; Farrar et al., 2001; Hawker et al., 2011). Because this score’s polarity is reversed compared to the other WB scores where a greater value reflects a negative WB, the score was reverse coded for consistency (i.e., greater scores reflected greater physical WB).
5) **Social WB** using the inclusion of the Other in Self (Aron et al., 1992, 1991), which captures how close and connected the respondent feels with other individuals or groups.

### EEG and ECG data collection

Once participants completed the survey, 2 minutes of EEG and ECG were recorded. Participants were instructed to close their eyes and focus on their breath by mentally counting each inhalation/exhalation cycle and to bring back their attention to their breath when they noticed that their minds were wandering. 4-channel EEG was recorded using Muse(version 2, 2016; InteraXon Inc.) with a 256 Hz sampling rate and 12-bit resolution. This system has 4 active electrodes (2 frontal and 2 temporoparietal ones), a passive-driven right leg, and an active common mode sense reference at the FPz location. It can be adjusted to different sizes (52-60 cm range). The skin under electrode sites was cleaned with alcohol swipes, and a thin layer of water was applied to the EEG electrodes with a sponge to improve signal quality.

The ECG electrode was custom-built to connect to the Muse’s auxiliary port following the manufacturers’ instructions. A 13-mm male nickel snap termination was connected to a shielded wire (part# ARF2168-ND @ Digikey) with 1/16” shrink tubing (for insulation). Then, the other end of the shielded wire was soldered to pin 4 of a micro-USB connector (part# H11613-ND @ Digikey) and sealed with hot glue and shrink tubing. This ECG electrode uses the same built-in Fpz reference as above for EEG. Disposable male snap sticky ECG electrodes with electroconductive gel (commonly available bio-medical.com) were connected to the Muse and easily attached to the participant’s wrist.

Simultaneous EEG and ECG signals were streamed with Bluetooth low energy to the MindMonitor App (Clutterbuck, 2015) on a Chromebook and automatically exported as .csv files into a Dropbox at the end of the recording. The MindMonitor App provided a basic electrode contact confirmation and signal quality was assessed visually by an EEG specialist using the raw signal displayed on the screen in real-time.

### Signal processing

CSV files were imported into EEGLAB v2023 (Delorme and Makeig, 2004) in Matlab R2023b (The MathWorks, Inc.) using the *import_muse* function (Cannard, 2021). The ECG signal was separated from the EEG signals for their subsequent processing.

#### EEG

3D electrode coordinates were imported using the boundary element model (BEM), and EEG signals were bandpass-filtered at 1-50 Hz using *pop_eegfiltnew()* (noncausal linear-phase FIR filter; order = 846; transition bandwidth = 1 Hz). Since traditional methods to automatically detect abnormal EEG channels cannot be used with only 4 electrodes, a machine learning (ML) classifier method was used (described in more detail in Cannard et al., in review). In brief, a selection of features is extracted from EEG signal on 5-s sliding windows and the trained models classify each window as good/bad ( 93.5% accuracy for the frontal channels and 91.4% accuracy for the posterior channels). The whole channel was removed if 50% of the windows were flagged as bad or contained a flat line. On average, 0.2 channels were removed per subject (SD = 0.5). These channels could not be interpolated because of the low number of electrodes available. EEG artifacts (hardware- or subject-related) were rejected automatically using the artifact subspace reconstruction method (ASR; *clean_asr()* function; (Kothe and Jung, 2016; Miyakoshi, 2023; Mullen et al., 2015) with the validated cutoff of 11 (Delorme and Martin, 2021). On average, 22.4 s of EEG signal was removed per subject (SD = 18.5). Finally, the following EEG features were extracted using the BrainBeats EEGLAB plugin v1.4. (Cannard et al., 2023): individual alpha frequency (IAF), frontal and posterior alpha asymmetry when both electrode pairs were available, signal fuzzy entropy (similar to the conventional sample entropy measure of signal complexity, but more sensitive and robust with noisy signals; Azami et al., 2019).

#### ECG

ECG signal was processed using the BrainBeats EEGLAB plugin v1.4. (Cannard et al., 2023). This included bandpass filtering, extraction of the RR intervals (i.e., time elapsed between two successive R-peaks of the QRS complex), detection and interpolation of the RR artifacts (e.g., electrode disconnection, atrial ectopic beats, arrhythmias, etc.), obtaining NN intervals (i.e., normal to normal), and calculation of the validated Signal Quality Index (SQI; Vest et al., 2018). Following guidelines, files with at least 20% of the signal with an SQI below 0.9 were considered “bad” and excluded from the analysis. The following HRV metrics were extracted from the NN intervals using the BrainBeats EEGLAB plugin: the standard deviation of the NN intervals (SDNN; time domain), the normalized low-frequency to high-frequency power ratio (LF/HF ratio; frequency domain), the short-to-long term NN interval standard deviation (Poincare SD1/SD2 ratio; nonlinear domain), fuzzy entropy (nonlinear domain similar to sample entropy but more robust; Azami et al., 2019).

### Statistics

Statistical analyses were performed using the Robust Correlation toolbox (Pernet et al., 2013) in MATLAB 2023b. All features (WB, HRV, and EEG) were gathered into a correlation matrix. Pairwise correlations were performed for each possible pair of variables using skipped Spearman’s correlations, which account for non-normal distributions, heteroscedasticity, and bivariate outliers (Pernet et al., 2013; Rousselet and Pernet, 2012; Wilcox et al., 2018). The resulting *p-values* were corrected for multiple comparisons (type 1 or family-wise error) using the Hochberg method (Pernet et al., 2013; Rousselet and Pernet, 2012; Wilcox et al., 2018). Only the significant correlations are reported in a correlation matrix, with circle size indicating the strength correlation coefficients and colors indicating the valence (warm colors for positive correlation and cold colors for negative correlations). The upper triangle was removed as these data are symmetric.

## RESULTS

### Signal Quality

Of the 60 initial participants, 8 were removed because the ECG signal quality index was too low (see Methods; Vest et al., 2018). EEG data quality was satisfactory, with an average of 0.2 channels ( *SD = 0.5*) removed per participant and an average of 22.4 s (*SD = 18.5*) of artifacts removed per participant. In the remaining files, 67.3% of participants were women; participants’ ages ranged from 22 to 73 (*M = 53; SD = 13*).

### Well-Being, EEG, and HRV

A correlation matrix (**Figure 1**) summarizes the pairwise Spearman correlations that were significant after correction for multiple comparisons (type 1 error).

**Figure 1.**
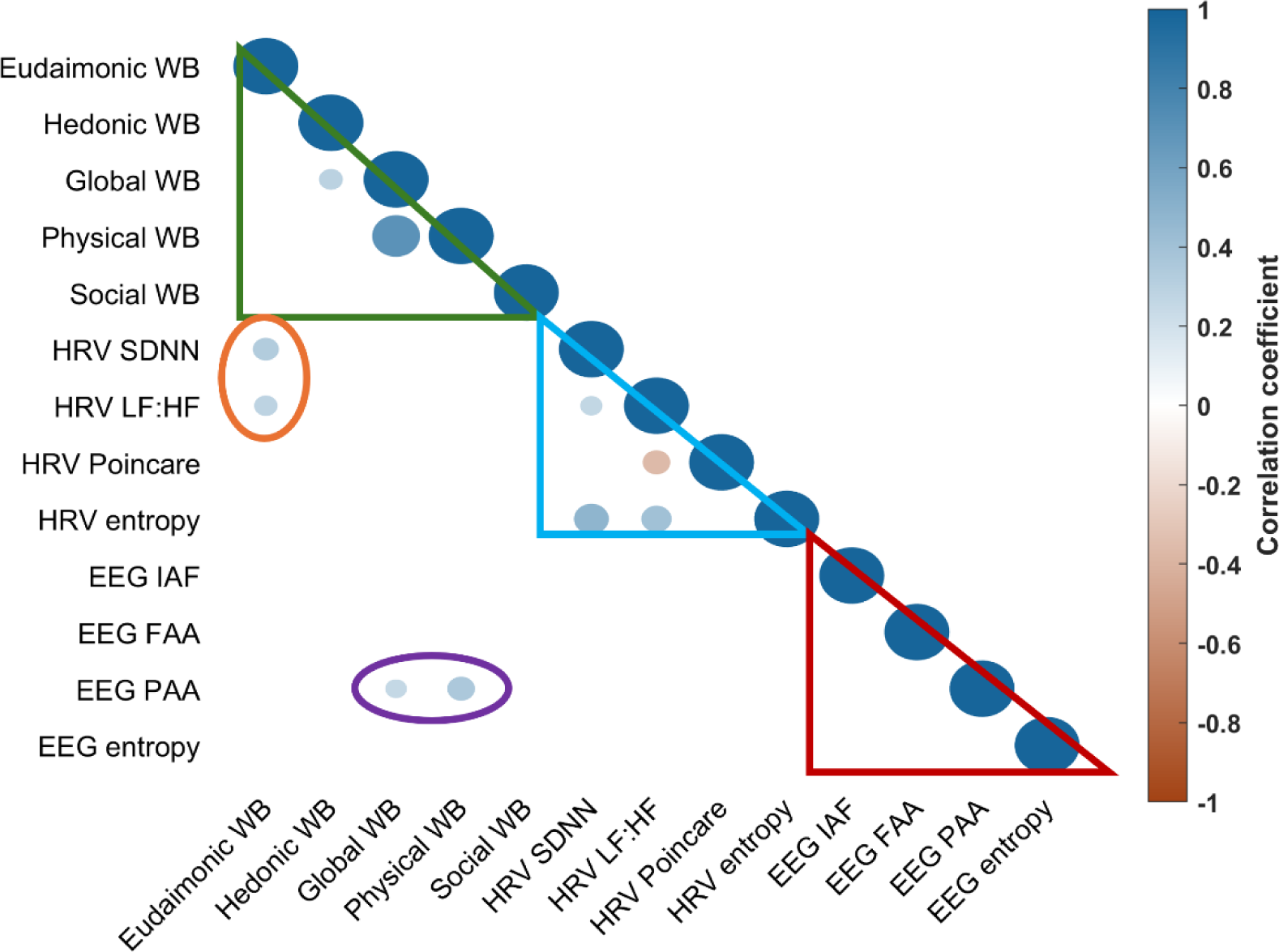
Correlation matrix of the pairwise correlations. Only the correlations significant after correction for multiple comparisons (family-wise error) at the 95% confidence level (p<0.05) are shown. The size of the circles reflects the correlation coefficients. Cold colors represent positive correlations, whereas warm colors represent negative correlations. The green triangle shows correlations between WB dimensions. The blue triangle shows correlations between HRV metrics. The red triangle shows correlations between EEG metrics. The orange circle shows correlations between HRV and WB measures. The purple circle shows correlations between EEG and WB measures. Significant correlations are reported in detail in the Results section.

As expected, the WB measures correlated with each other. Global WB was weakly correlated with Hedonic WB (*Rho = .3; p-corrected= 0.034*) and strongly correlated with physical WB (*Rho = .7; p-corrected = 0.001*).

Eudaimonic WB was positively correlated with two HRV metrics, SDNN ( *Rho = .34; p-corrected= 0.018*) and the LF/HF ratio (*Rho = .29; p-corrected= 0.022*). Furthermore, as reported by previous investigators (Shaffer et al., 2020; Shaffer and Ginsberg, 2017), several HRV metrics were correlated (e.g., LF/HF ratio, SDNN, entropy, positively, and the Poincare SD1/SD2 ratio negatively).

EEG posterior alpha asymmetry was positively correlated with global WB ( *Rho = .26; p-corrected = 0.024*) and Physical WB (*Rho = .36; p-corrected = 0.044*).

No relationships were observed between WB and the IAF, the frontal alpha asymmetry, and nonlinear metrics (HRV Poincare, HRV entropy, and EEG entropy). In addition, no associations were found between the EEG and HRV measures.

## Discussion

### Results summary

#### Signal quality

Fifty-two out of sixty ECG files had a good signal quality index (See Methods). This is better than we expected, considering the low cost and portability nature of the recording hardware and the location of the ECG electrodes on the participants’ wrists. With 5% of channels and 19% of the signal removed on average per participant, EEG signal quality was also estimated to be satisfactory according to our quality criteria (see Methods). While the percentage of artifacts or artifact-free data is often considered a signal quality index, future studies could develop a standardized, universal signal quality index for low-density wearable EEG (e.g., (Gabard-Durnam et al., 2018; Kaiser et al., 2021; Lucey et al., 2016; Mahdid et al., 2020).

#### WB & HRV

HRV SDNN and LF/HF ratios were positively associated with Eudaimonic WB, in line with some previous findings (Appelhans and Luecken, 2006; Shaffer et al., 2014). Eudaimonic WB entails the sense of making personal sacrifices for a greater cause and the sense of alignment between natural responses and long-term goals. Individuals who engage in activities that they perceive as contributing to the greater good tend to report higher levels of life satisfaction. The sense of purpose and commitment to something larger than oneself is considered a core element of psychological WB, as it transcends mere personal gratification and involves striving toward meaningful goals. Additionally, living in a way consistent with one’s true self, personal values, and long-term aspirations fosters authenticity and greater psychological WB. When individuals experience congruence in their lives, they are more likely to engage in personally fulfilling activities that align with their broader life goals, enhancing their eudaimonic WB. Consequently, higher scores in this WB dimension may be associated with increased ANS balance.

#### WB & EEG

We observed a significant relationship between posterior alpha asymmetry, Global WB, and physical WB. These findings conform to previous findings, confirming the relationship between posterior alpha asymmetry and global WB (Cannard et al., 2021b). Interestingly, the correlation between Global and Physical WB suggests that the Global WB scores were dominantly modulated by participants’ perceived sleep disturbance and pain intensity levels. Hence, these findings indicate that posterior alpha asymmetry may reflect brain processes that regulate physical discomfort resulting from pain or fatigue.

#### EEG and HRV

While some previous studies found associations between EEG and HRV signals during sleep (Abdullah et al., 2010; Ako et al., 2003), we did not find any association between these two biosignals in this study. Interestingly, these previous studies found correlations between EEG delta and LF-HRV power (and the LF/HF ratio) during sleep. Delta power is known to be associated with the glymphatic system (facilitates the clearance of waste and soluble proteins from the central nervous system), which involves high vascular (e.g., the opening of the blood-brain barrier) and cerebrospinal activity. Thus, these previous associations may be specific to sleep rather than waking states, as in our study.

### Limitations

Certain limitations may affect the interpretation of these results. First, the study involved secondary analyses of a previously collected dataset. Therefore, future studies should aim to repeat these analyses as primary research questions incorporating optimal study design to answer those questions. We aimed to assess whether one could rapidly but reliably assess WB measures using short and easily collected subjective questionnaires and objective physiological tools. However, while these questionnaires were validated, future studies may benefit from using more lengthy multi-item questionnaires to capture the complex subjective WB constructs and, therefore, identify stronger relationships with physiological measures. Similarly, EEG and HRV metrics are more reliable when estimated on longer measurements (e.g., 15 minutes). Hence, investigators may identify stronger relationships between HRV/EEG and WB using more robust estimates, which can be used in later studies to quantify WB using portable, ultrashort physiological measurements alone.

The task during which the ECG and EEG signals were collected was a breath-counting exercise. Shifts in respiration rate and volume can influence HRV measures, which do not necessarily affect vagal tone (Shaffer and Ginsberg, 2017). Thus, individuals with greater eudaimonic WB may have developed emotion regulation or other coping strategies that involve modulations of their respiration to successfully fulfill their goals when faced with stressors (i.e., indicating resilience).

The ECG electrodes show good signal quality. However, they are not commercially available and need to be constructed. While their construction is relatively easy and can be implemented with instructions provided by the InteraXon website, making them commercially available would greatly facilitate this research of related applications.

### Future directions

We selected theory-driven EEG and HRV metrics that seemed promising for the goals of our study. To avoid excessive numbers of comparisons and increasing the type 1 error, we kept the number of metrics to a reasonable number. However, we confirmed previous findings showing that HRV metrics are inter-correlated. Also, some variables did not present any relationship with WB, suggesting they could be removed in future analyses. Future studies investigating WB may, therefore, reduce HRV metrics to the LF/HF ratio only, which showed the strongest association, in addition to posterior alpha asymmetry.

Contrary to expectations, nonlinear measures (Poincare HRV, entropy) did not show significant relationships. Despite these findings, more research is needed on these and other new metrics associated with the psychological and physical components of well-being. One such metric is frontal theta “cordance” (i.e., the proportion of absolute theta power relative to the total power), which is associated with predicting the response to antidepressant treatment (e.g., Baskaran et al., 2012; Olbrich and Arns, 2013). Similarly, frontal beta power may potentially be associated with WB, as it is associated with stress, anxiety, and depression (e.g., Hamid et al., 2010; Jun and Smitha, 2016). The theta/beta ratio may also be a promising indicator of stress, resilience, and ruminations, and therefore WB (Putman, 2011; Putman et al., 2014; van Son et al., 2019).

Larger samples are required to confirm these findings. The tools and methods in this study should allow future investigators to collect large and diverse sample sizes necessary to leverage machine learning (ML) analyses that will increase the accuracy of these potential biomarkers of WB and help to forecast and classify WB status in real-world settings. While we did not find associations between EEG and HRV measures, future researchers may combine features from these two types of biosignals for more performant ML models (Ahn et al., 2019).

## Conclusion

We report the feasibility of using low-cost, portable tools to measure relevant ultrashort EEG and ECG signals. The resulting HRV metrics obtained from these ECG signals may be useful for capturing information about autonomous nervous system activity associated with psychological WB (sense of purpose, fulfillment, alignment). In contrast, EEG metrics may capture cognitive activity related to the physical dimension of WB (sleep and pain levels).

## Acknowledgment

We wish to thank the Institute of Noetic Sciences, the Region Occitanie, France (LSP Grant #183258, BIGDAT, UMR5549) for funding this work, as well as Jim & Christina Grote, the Biofield Tuning Institute, the John Brockway Huntington Foundation, Thomas Matte, the Patricia Beck Phillips Foundation, and the John Sperling Foundation for supporting this work. We would also like to thank Kenneth Rachlin for building the ECG electrodes used in this study.

